# Genome assembly of the common pheasant *Phasianus colchicus*, a model for speciation and ecological genomics

**DOI:** 10.1101/818666

**Authors:** Yang Liu, Simin Liu, Nan Zhang, De Chen, Pinjia Que, Naijia Liu, Jacob Höglund, Zhengwang Zhang, Biao Wang

## Abstract

The common pheasant (*Phasianus colchicus*) in the order Galliformes and the family Phasianidae, has 30 subspecies distributed across its native range in the Palearctic realm and has been introduced to Europe, North America, and Australia. It is an important game bird often subjected to wildlife management as well as a model species to study speciation, biogeography and local adaptation. However, the genomic resources for the common pheasant are generally lacking. We sequenced a male individual of the subspecies *torquatus* of the common pheasant with the Illumina Hiseq platform. We obtained 94.88 Gb of usable sequences by filtering out low-quality reads of the raw data generated. This resulted in a 1.02 Gb final assembly, which equals the estimated genome size. BUSCO analysis using chicken as a model showed that 93.3% of genes were complete. The contig N50 and scaffold N50 sizes were 178 kb and 10.2 Mb, respectively. All these indicate that we obtained a high-quality genome assembly. We annotated 16,485 protein-coding genes and 123.3 Mb (12.05 % of the genome) of repetitive sequences by ab initio and homology-based prediction. Furthermore, we applied a RAD-sequencing approach for another 45 individuals of seven representative subspecies in China and identified 4,376,351 novel single nucleotide polymorphism (SNPs) markers. Using this unprecedented dataset, we uncover the geographic population structure and genetic introgression among common pheasants in China. Our results provide the first high-quality reference genome for the common pheasant and a valuable genome-wide SNP database for studying population genomics and demographic history.

## Introduction

The common pheasant (*Phasianus colchicus*), belonging to the order Galliformes in the family Phasianidae, is a common gamebird with a worldwide distribution (Hill & Robertson, 1988; Pfarr, 2012). It is a well-known game bird with global importance for both research and wildlife management. First, as one of the world’s most widespread resident species, the common pheasant is native to the temperate zones in the Palearctic region, from far eastern Siberia to eastern-southeastern Europe (east of the Black Sea), and southwards to Indochina and Afghanistan (Cramp, 1980; Johnsgard, 1999). Second, it is among the most subspecies-rich bird species with thirty described subspecies defined mainly by plumage characters in males and geographical range (Madge, McGowan, & Kirwan, 2002). These subspecies occupy substantially different environments and climatic zones, from isolated oases in semi-deserts, to montane regions, displaying unique phenotypes and genotypes (Hill & Robertson, 1988; Pfarr, 2012). The common pheasant also has great economic values, and have long history of being released for hunting or kept captive in bird farms in Western Europe, North America and Australia (Johnsgard, 1999; Madge et al., 2002). This makes the common pheasant a promising model to investigate important questions about speciation, trait evolution, biogeography and local adaptation to various climatic conditions, as well as in wildlife management and conservation genetics.

Previous studies of the common pheasant have mainly focused on its ecology and biology. Several aspects of the species’ life history and reproduction, such as survival rate (Draycott, Hoodless, Woodburn, & Sage, 2008), habitat selection (Long, Zhou, Wang, Wei, & Hu, 2007) and breeding biology (Robertson, 1996) has been well quantified, which provide useful knowledge for sustainable management. The taxonomy and systematics of common pheasant have long been actively studied. For example, the thirty-recognized subspecies were hypothesized to be clustered into five subspecies-groups based on morphological resemblance and biogeographical affinity (Madge et al., 2002). However, given that clinal variation may explain connectivity and contiguous distributions, the validity of some subspecies have been questioned (Cramp, 1980; Johnsgard, 1999; N. Liu & Sun, 1992; Madge et al., 2002). Molecular phylogenetic approaches have been applied to resolve these puzzles since the subspecies-groups have been identified as independent evolutionary lineages by mitochondrial fragments (Y. Liu, Zhan, Wang, Chang, & Zhang, 2010; Qu, Liu, Bao, & Wang, 2009; Zhang, An, Backström, & Liu, 2014). Recent work using multi-locus nuclear markers corroborate previous findings, and further showed that viscous boundaries between subspecies are probably due to extensive gene flow among contiguous populations (Kayvanfar, Aliabadian, Niu, Zhang, & Liu, 2017; Wang et al., 2017). This work used a small number of genetic markers, which is capable of delineating population structure and subspecies relationship. However, due to the lack of genomic-level data, many questions related to demographic histories (e.g. (Nadachowska-Brzyska et al., 2013)), population admixture and molecular genetic basis of phenotypes (e.g. (Toews et al., 2016)), remain undetermined.

Taking advantage of next-generation sequencing (NGS), it is now feasible to obtain unprecedented genomic-scale data, allowing direct quantification of genomic variation and identification of the genomic landscape associated with specific phenotypes in non-model organisms (Bosse et al., 2017; Ellegren, 2014; Toews et al., 2016). Particularly in the study of speciation, it is of interest to understand the size and extent of regions of divergence across the genome among the related species (Wu, 2001). Recent empirical studies have been uncovering a notable pattern that genomic regions with accumulated genetic differentiation between closely related avian species/subspecies pairs contain genes involved in divergence while gene flow and genetic drift homogenize other regions (Ellegren et al., 2012; Poelstra et al., 2014). Divergent heterogeneous regions have been coined “islands of divergence” (Harr, 2006; Nosil, Funk, & ORTIZ-BARRIENTOS, 2009; Turner, Hahn, & Nuzhdin, 2005). Although strong divergent selection is considered to contribute to such genomic islands, the underlying evolutionary processes driving this notable genome-wide pattern is still controversial. Other confounding evolutionary processes, such as genetic drift, may also contribute. Given the common pheasant is distributed in a vast geographical range with varied environment conditions and habitats, one can predict that complex evolutionary processes and demographic history could contribute to population divergence among populations of the common pheasant. Therefore, it is extremely interesting to screen the genomic landscape affected by the aforementioned factors. Obviously, the availability of genomic information will help facilitate finer-scale characterization of the genomic-wide pattern of population divergence.

In this study, we present the first genome assembly of the common pheasant with detailed description of its genetic architecture and population-level genomic polymorphisms in order to provide genomic resources towards studies of speciation, local adaptation and conservation genomics of an ecologically important species.

## Materials and Methods

### Sampling and Sequencing

The sequenced sample was fresh blood from a male common pheasant, (the subspecies *Phasianus colchicus torquatus*: NCBI taxonomy ID 9054; BioProject ID PRJNA449162; BioSample ID SAMN08888528). This individual was a captive individual with a wild origin from Beijing Wild Animal Park (Daxing) sampled in August 2015 (Figure 1). All sampling collection was performed in accordance with Chinese wildlife regulations and protocols.

**Fig. 1.**
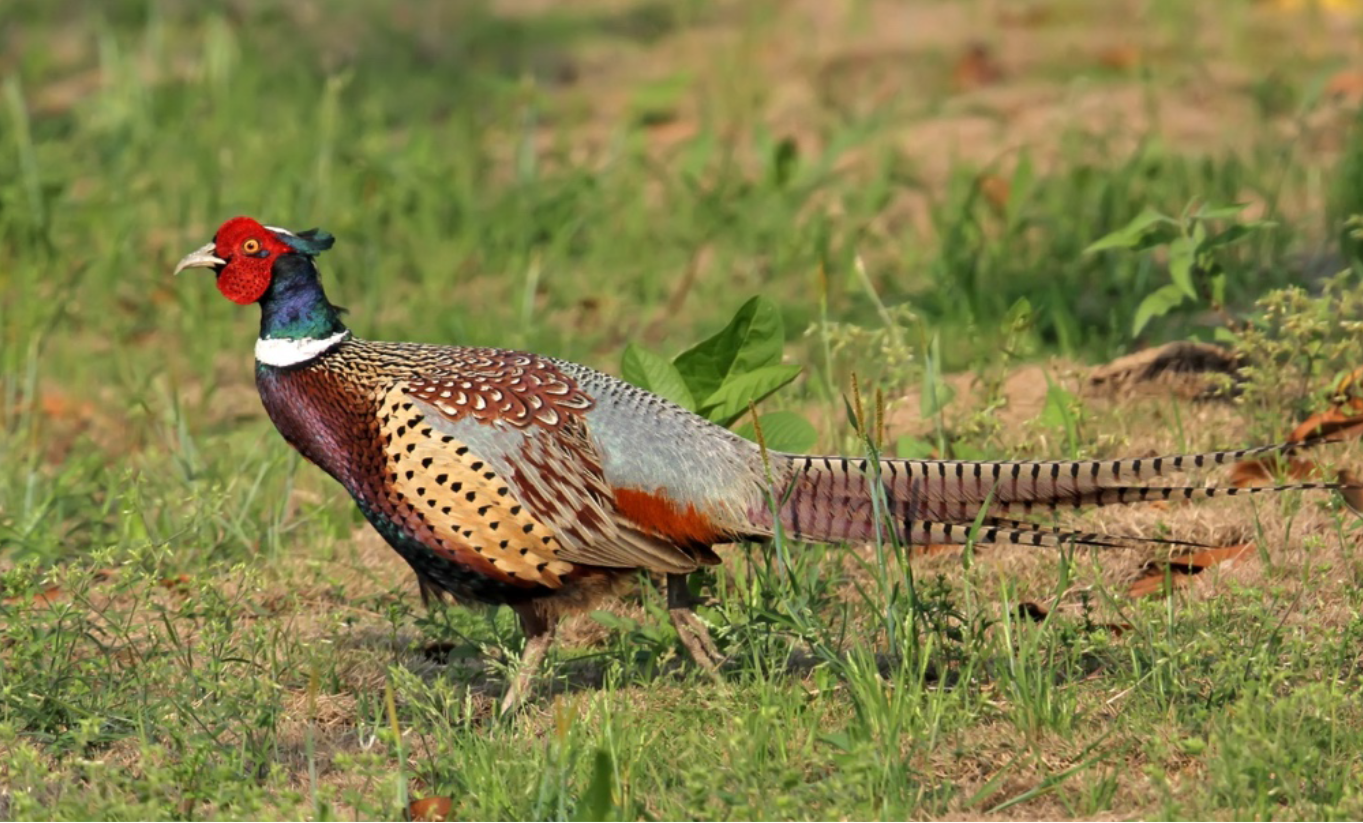
The common pheasant *Phasianus colchicus*, where the subspecies *torquatus* was sequenced in this study.

Genomic DNA was extracted using a Qiagen DNA purification kit following the manufacturer’s instruction and the quality of extracted DNA was checked using gel electrophoresis (1% agarose gel/40ng loading). We built four short insert libraries (two for 250 bp and two for 450 bp) and three mate pair libraries (2kbp, 5kbp, 10kbp) following Illumina’s standard protocol. Briefly, the qualified genomic DNA was randomly sheared into short fragments by hydrodynamic shearing system (Covaris, Massachusetts, USA). Then, followed by end repairing, dA-tailing and further ligation with Illumina adapter, the required fragments (in 300-500 bp size) with both P5 and P7 sequences were PCR selected and amplified. After gel electrophoresis and subsequent purification, the required fragments were obtained. The constructed libraries were loaded on the Illumina HiSeq platform for paired-end sequencing, with the read length of 150 bp at each end. Raw data obtained from sequencing also contains adapter contamination and low-quality reads. These sequence artifacts may complicate the downstream processing analysis. The raw data was thus filtered to reduce low quality bases and reads by the following strategies: (i) filtering our reads with adapters; (ii) reads with N bases more than 5%; (iii) the paired reads when single end sequencing reads contain low quality (<5) bases that exceed 10% of the read length.

### Evaluation of genome size

The genome size was estimated according to a k-mer analysis with the formula: G = k-mer_number / k-mer_depth, where G is the genome size, k-mer_number is the total counts of kmers and k-mer_depth refers to the main peak in the k-mer distribution. In this study, we collected all reads in a short-insert library to conduct the 19-mer analysis with Jellyfish 2.0 (Marçais & Kingsford, 2011). A total of 39,151,211,661 k-mers were produced and the peak k-mer depth was 38 (Figure S1).

### Genome assembly and assessment

To assemble the common pheasant genome, we firstly evaluated genome-wide heterozygosity using the above k-mer analysis. Double peaks suggested that this diploid genome was highly heterozygous. We therefore employed Platanus v1.2.4 (Kajitani et al., 2014), which is particularly designed for highly heterozygous genomes, to assemble the common pheasant genome. The first round included three steps: contig assembly, scaffolding, and gap closing. Firstly, all filtered reads in short-insert libraries (250bp, 450bp) were input for contig assembly. After constructing de Brujin graphs, clipping tips, merging bubbles, and removing low coverage links with default parameters, assembled contigs and bubbles in the graphs were obtained in this step. In the scaffolding steps, the bubbles and reads from both short-insert library (250bp, 450bp) and long-insert library (2kbp, 5kbp,10kbp) were mapped onto contig sequences to build scaffolds with default parameters. Finally, the intra-scaffold gaps were filled with reads from all libraries. After five rounds of gap closing, the gap rate in the scaffolds reached a plateau. To evaluate the completeness of the genome, we performed BUSCO v3 (Simão, Waterhouse, Ioannidis, Kriventseva, & Zdobnov, 2015) with the representative chicken gene dataset aves_odb9.

### Genome annotation

To identify genomic repeat elements in the assembly, both ab initio and homolog-based methods were used. For the homolog-based methods, we used RepeatMasker (www.repeatmasker.org) (Smit, Hubley, & Green, 2016) to search against the Repbase library version 22.12 (Jurka, 1998). In the ab initio method, a custom repeat library was constructed using RepeatModeler (www.repeatmasker.org) (Smit & Hubley, 2008) with RECON (Bao & Eddy, 2002), RepeatScout (Price, Jones, & Pevzner, 2005) and Tandem repeats finder (TRF) (Benson, 1999), which was then used in RepeatMasker to annotate repeats.

Gene annotation for the common pheasant genome assembly were conducted with the MAKER2 pipeline (Holt & Yandell, 2011), which incorporates ab initio prediction and homology-based prediction. For the ab initio method, repeat regions were first masked based on the previous results of repeat annotation, and then Augustus (Stanke & Waack, 2003) and GeneMark_ES (Lomsadze, Ter-Hovhannisyan, Chernoff, & Borodovsky, 2005) were employed to generate gene structures. In addition, FGENESH (Salamov & Solovyev, 2000) was also used to for ab initio prediction. For homology-based prediction, protein sequences from three different species, Chicken (*Gallus gallus*), Zebra finch (*Taeniopygia guttata*), Turkey (*Meleagris gallopavo*) (downloaded from Ensemble database 9.1 release), were mapped onto the genome assembly using tBlastn of the NCBI BLAST suite v2.7.1 (Coordinators, 2017; Madden, 2013) and Exonerate v2.2.0 (Slater & Birney, 2005) was used to polish BLAST hits to get exact intron/exon position.

### Population genomics analysis using the reference genome of the common pheasant

To facilitate future population genomics studies of the common pheasant, we further sequenced an additional 45 male individuals from seven subspecies across China using restriction site-associated DNA sequencing (RAD sequencing) (Miller, Dunham, Amores, Cresko, & Johnson, 2007) (Figure 2a, Table S4). The de novo common pheasant genome was used as a reference to facilitate mapping and SNP calling. We analyzed population structure among 45 individuals using ADMIXTURE 1.3 (Alexander, Novembre, & Lange, 2009) and principal component analysis (PCA). The detailed lab and sequencing procedures, and analyses were available in the Appendix of Supplementary Materials.

## Results and Discussion

We sequenced and assembled a reference genome of a male common pheasant. We obtained 94.88Gb clean paired reads (Table S1). The assembled genome size of is 1.02Gb (1,021,360,992bp) in length with a genomic coverage =93x. The assembled genome contains 58,369 contigs (contig N50 of 178kb) and 39,677 scaffolds (scaffold N50 of 10.2Mb) (Table S2). The completeness of the common pheasant draft genome is high: we totally identified 4790 BUSCOs (97.5%) including 4,585 complete (93.3%) and 205 fragmented (4.2%) BUSCOs (Table S3).

We found that a total of 12.05% (123.3 Mb) repeats elements were identified in the genome assembly of common pheasant, with unclassified elements constituting the greatest proportion (Table 1). We used different prediction methods to produce a consensus gene set. In total 16,485 protein-coding genes were identified in the common pheasant genome using our described prediction methods (Table 2).

**Table 1.** Summary of identified repeat elements in the genome assembly of the common pheasant.

|  | Number of elements | Length occupied | Percentage of sequence |
| --- | --- | --- | --- |
| SINEs | 330 | 91,680 bp | 0.01 % |
| LINEs | 81179 | 15,470,955 bp | 1.51 % |
| LTR elements: | 15921 | 4,853,455 bp | 0.47 % |
| DNA elements | 7847 | 2,561,151 bp | 0.25 % |
| Unclassified | 279128 | 82,169,029 bp | 8.03 % |
| Small RNA | 273 | 86,874 bp | 0.01 % |
| Satellites | 8976 | 3,506,208 bp | 0.34 % |
| Simple repeats | 307238 | 12,025,779 bp | 1.18 % |
| Low complexity | 55326 | 2,838,076 bp | 0.28 % |
| **Total** |  | 123,323,121 bp | 12.05 % |

**Table 2.**
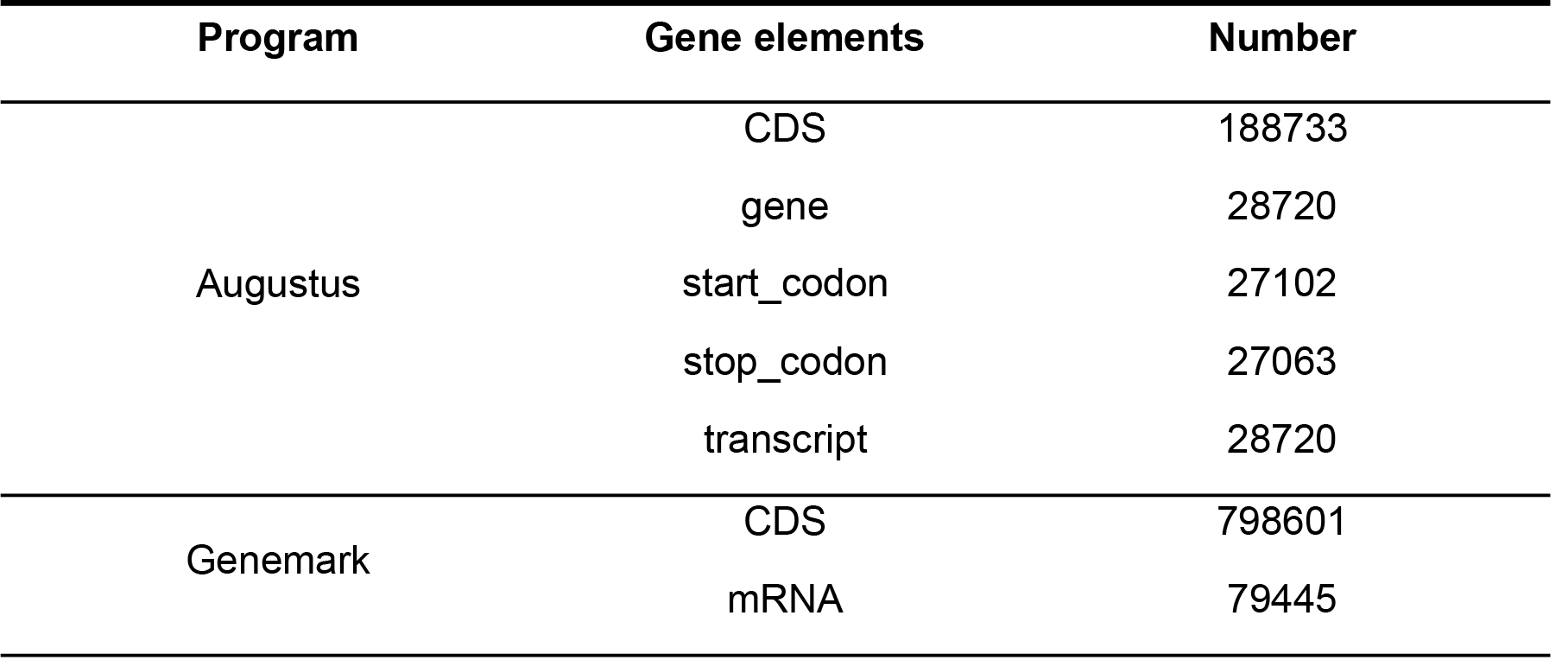
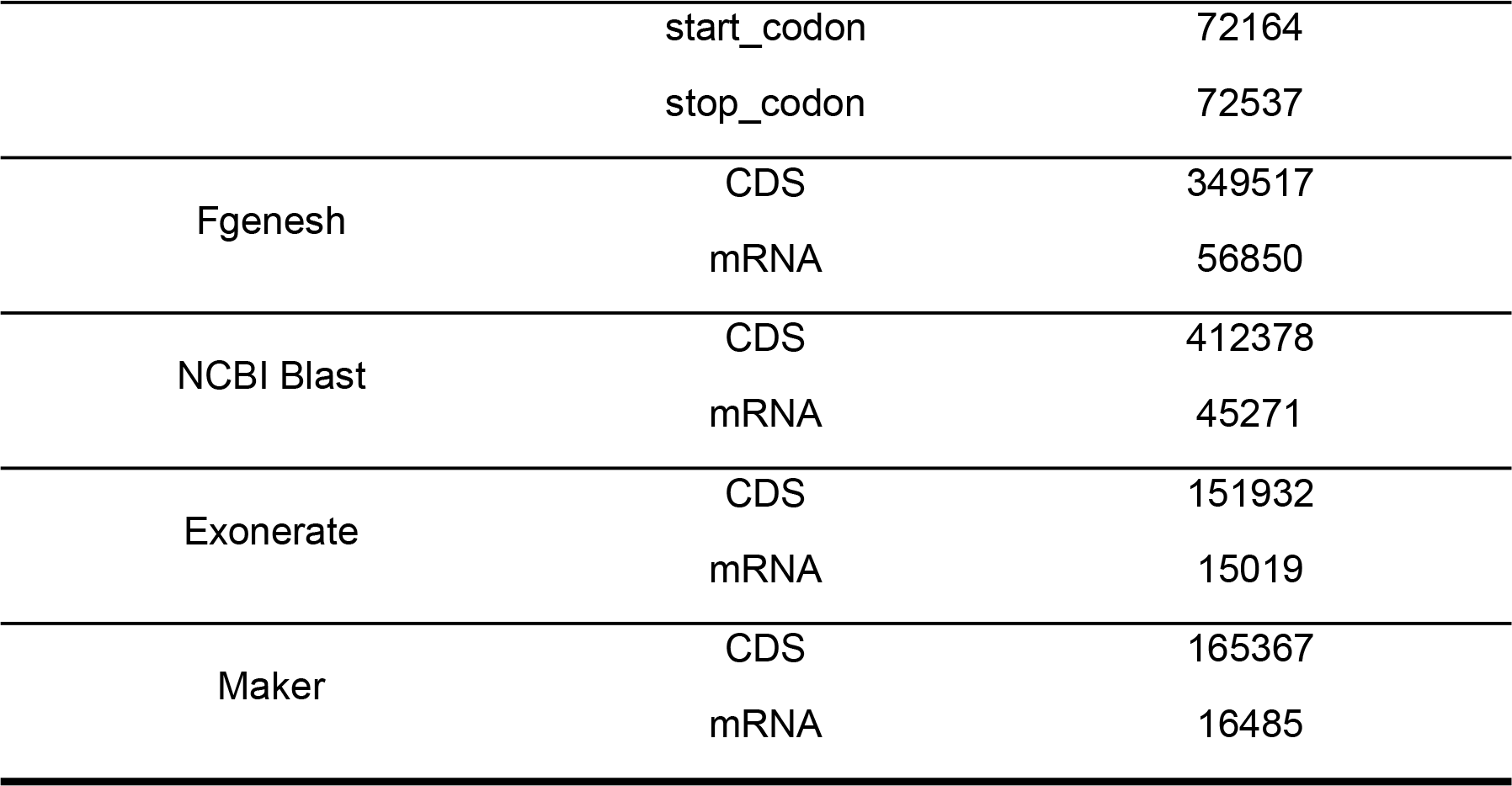
Gene annotation of the common pheasant genome.

| Program | Gene elements | Number |
| --- | --- | --- |
| Augustus | CDS | 188733 |
|  | gene | 28720 |
|  | start_codon | 27102 |
|  | stop_codon | 27063 |
|  | transcript | 28720 |
| Genemark | CDS | 798601 |
|  | mRNA | 79445 |
|  | start_codon | 72164 |
|  | stop_codon | 72537 |
| Fgenesh | CDS | 349517 |
|  | mRNA | 56850 |
| NCBI Blast | CDS | 412378 |
|  | mRNA | 45271 |
| Exonerate | CDS | 151932 |
|  | mRNA | 15019 |
| Maker | CDS | 165367 |
|  | mRNA | 16485 |

In addition, about 114.91 Gb were generated (clean data) for all 45 samples. Data summary is shown in Table S5. After filtering and SNP callings, we obtained 4,376,351 novel SNPs overall, and we established a database including 328,473 to 999,733 SNPs for each individual (Table S4). We managed to identify 59,453 SNPs in exon regions, UTR regions and splice sites. 602,747 SNPs were identified in 5kbp upstream/downstream regions, which can also be associated with phenotypes and functions. In addition, 1,260,753 SNPs and 3,092,463 SNPs were found in intron regions and intergenic regions respectively. Intron regions are suggested to have genomic functions (Cech, 1990) and we can further test this hypothesis using the common pheasant model. Since populations of common pheasants dwelling in contrasting environments and climatic zones, e.g. monsoon regions, basins in the Qinghai-Tibetan plateau, semi-arid zones, and deserts. These resources can be used to investigate population genomics and genomic architectures associated with local adaptation of the common pheasant in the future (Table S5).

Our population structure results by ADMIXTURE clearly show that when four groups were inferred, the cross-validation error has the lowest value among the alternatives (Figure S2). Given this, populations in western Xinjiang (subspecies *shawii*), Qinghai (subspecies *vlanglii*), Yunnan (subspecies *elegans*) and the remaining populations in central and eastern China (subspecies *strauchi, kiangsuensis, karpowi* and *torquatus*) form distinct genetic groups, respectively (Figure 2b). The results of principal component analysis using the similar SNP dataset were consistent with Admixture results, showing four genetic clusters, i.e. Yunnan, western Xinjiang Qinghai, and the remaining subspecies. Clearly, the subspecies of *karpowi*, *kiangsuensis* and *torquatus* show pattern of genomic admixture with the subspecies of *vlanglii* from Qinghai, which corroborates previous results as shown by multilocus phylogeography studies [13][14]. The resulting population structuring pattern is likely due to genetic drift of small and isolated populations in those regions (Figure 2a), but consistent with genetic introgression caused by population expansion in contiguous populations in northern China (Liu et al. 2010; Kayvanfar et al. 2017). Obviously the derived genomic-level polymorphism provide unprecedented power, which allows exclusive tests of specific hypotheses demographic history in common pheasant using model-based population genomic analysis [49].

**Fig. 2.**
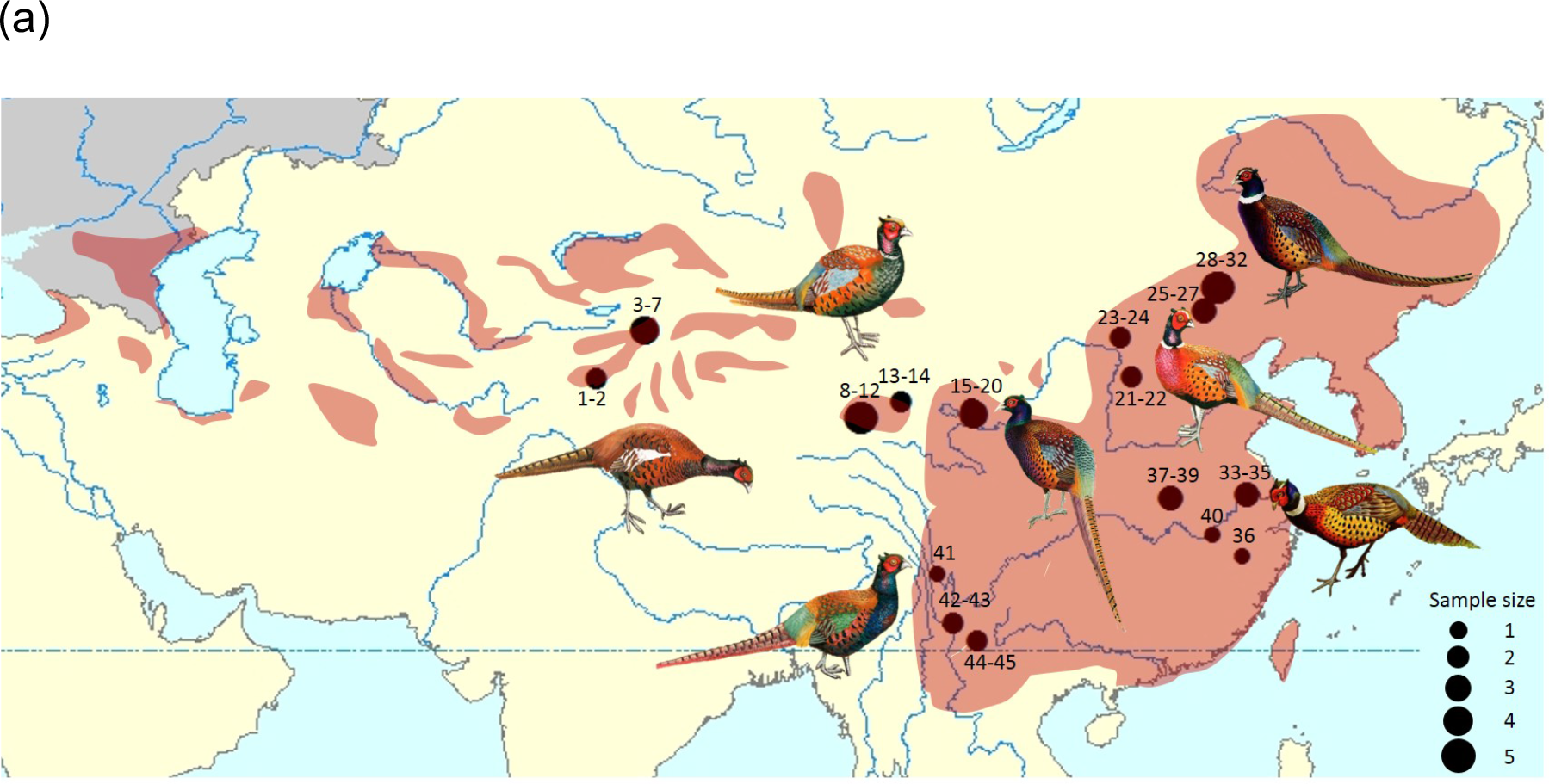
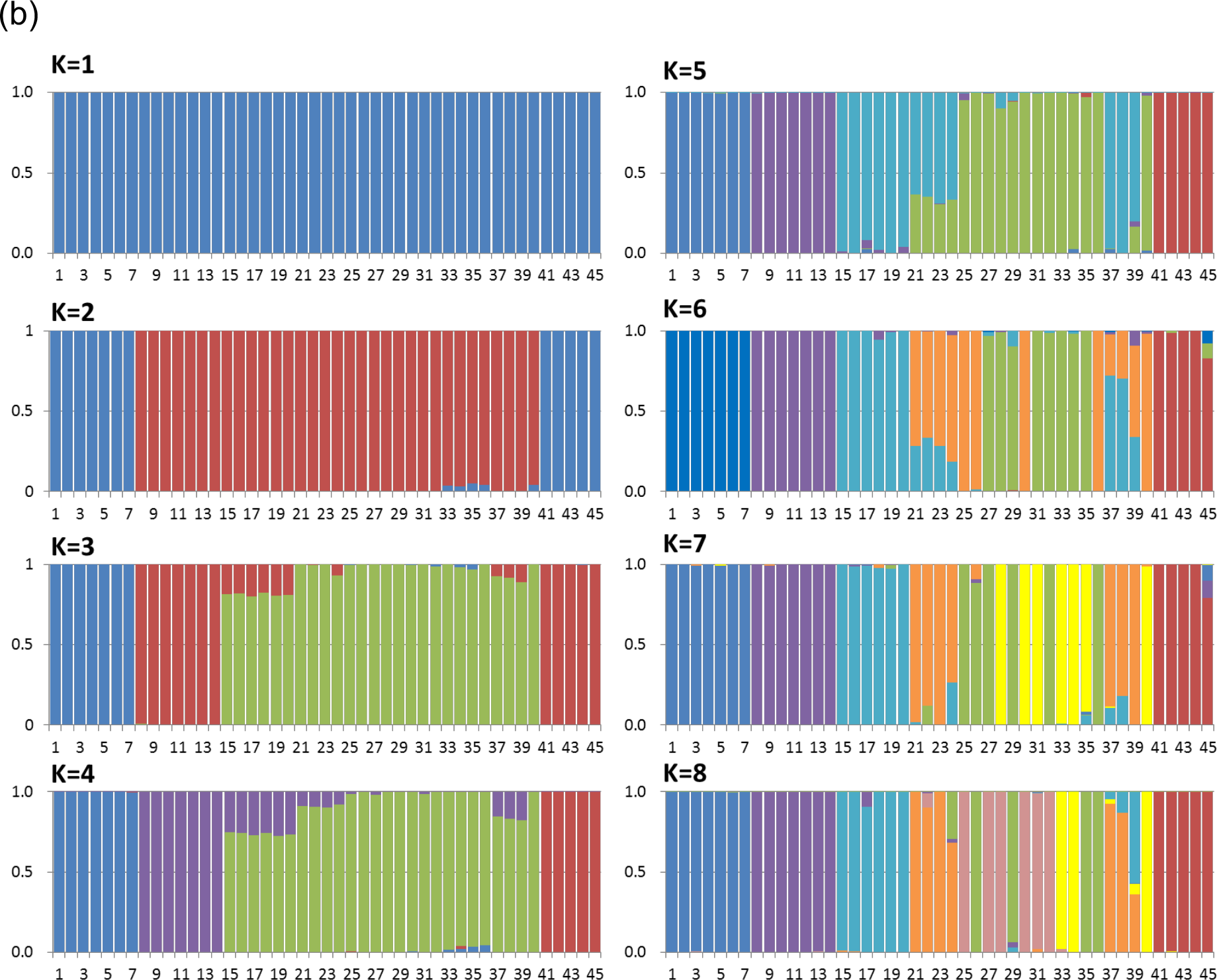
Population genetic structure of Common Pheasant *Phasianus colchicus* in China. (a) The distributions of samples of seven subspecies of common pheasant. Shown are males with divergent phenotypes (pheasant artworks were modified from Pfarr 2012). The size of the circles is proportional to the number of individuals. (b) Population structures of seven subspecies of the common pheasant in China. SNPs were derived using RAD-sequencing (K value 1~8, from upper left to lower right). The Y-axis represents proportion of each ancestral genetic component and numbers in the X-axis represent sample locations (details in Table S4) and subspecies affinities: 1-7: *shawii*; 8-14: *vlangalii*; 15-20: *strauchi*; 21-24: *kiangsuensis*; 25-32: *karpowi*; 33-40: *torquatus*; 41-45: *elegans*.

In conclusion, we report the first high quality assembled and annotated common pheasant genome. We also report a well-annotated SNP database of population samples for this species. These datasets provide valuable resources to the scientific community and open avenues to future deep investigation of questions in evolutionary biology such as speciation, trait evolution, biogeography, local adaptation and wildlife management in the common pheasant, a bird species of long-standing interest in ecology, evolution and economics.

## Acknowledgements

This work was supported by the National Natural Science Foundation of China (No. 3157225 to YL and No. 31702013 to BW). Partial computational machinery time work was granted by Special Program for Applied Research on Super Computation of the NSFC-Guangdong Joint Fund (2nd phase) under Grant No. U1501501 to YL. We also acknowledge support from Uppsala Multidisciplinary Center for Advanced Computational Science (SNIC-UPPMAX) for providing assistance in computational infrastructure. We thank the following persons who kindly provided samples or assisted with sampling: Xiaoju Niu, Ying Liu and Jun Gou. We thank Total Genomics Solution Institute, Shenzhen, China for bioinformatics advices.

## Author notes

### Data deposition

Raw sequencing reads are deposited in the NCBI Sequence Read Archive (SRR6987054 - SRR6987060), and the Whole Genome Shotgun project is deposited at NCBI GenBank under accession numbers BioProject PRJNA449162, BioSample SAMN08888240 and Genome assembly QCWP00000000. The SNPs are deposited at the European Bioinformatics Institute (EMBL-EBI) EVA database under accession number PRJEB26609.

## Literature Cited

Alexander, D. H., Novembre, J., & Lange, K. (2009). Fast model-based estimation of ancestry in unrelated individuals. Genome Research, 19 (9), 1655–1664.

Bao, Z., & Eddy, S. R. (2002). Automated de novo identification of repeat sequence families in sequenced genomes. Genome Research, 12 (8), 1269–1276.

Benson, G. (1999). Tandem repeats finder: a program to analyze DNA sequences. Nucleic Acids Research, 27 (2), 573.

Bosse, M., Spurgin, L. G., Laine, V. N., Cole, E. F., Firth, J. A., Gienapp, P., … Verhagen, I. (2017). Recent natural selection causes adaptive evolution of an avian polygenic trait. Science, 358 (6361), 365–368.

Cech, T. R. (1990). Conserved sequences and structures of group I introns: building an active site for RNA catalysis—a review. In RNA: Catalysis, Splicing, Evolution 300 (pp. 191–203): Elsevier.

Coordinators, N. R. (2017). Database resources of the national center for biotechnology information. Nucleic Acids Research, 45 (Database issue), D12.

Cramp, S. (1980). Birds of the western Palearctic: volume 2, hawks to bustards: Oxford University Press.

Draycott, R. A., Hoodless, A. N., Woodburn, M. I., & Sage, R. B. (2008). Nest predation of common pheasants Phasianus colchicus. Ibis, 150 (s1), 37–44.

Ellegren, H. (2014). Genome sequencing and population genomics in non-model organisms. Trends in ecology & evolution, 29 (1), 51–63.

Ellegren, H., Smeds, L., Burri, R., Olason, P. I., Backström, N., Kawakami, T., … Qvarnström, A. (2012). The genomic landscape of species divergence in Ficedula flycatchers. Nature, 491 (7426), 756.

Harr, B. (2006). Genomic islands of differentiation between house mouse subspecies. Genome Research, 16 (6), 730–737.

Hill, D., & Robertson, P. (1988). The pheasant: ecology, management and conservation: BSP Professional Books.

Holt, C., & Yandell, M. (2011). MAKER2: an annotation pipeline and genome-database management too tool for second-generation genome projects. BMC Bioinformatics, 12 (1), 491.

Johnsgard, P. (1999). The pheasant of the world: biology and natural history. Washington, DC: Smithsonian Institute, 2, 179–193.

Jurka, J. (1998). Repeats in genomic DNA: mining and meaning. Current Opinion in Structural Biology, 8 (3), 333–337.

Kajitani, R., Toshimoto, K., Noguchi, H., Toyoda, A., Ogura, Y., Okuno, M., … Maruyama, H. (2014). Efficient de novo assembly of highly heterozygous genomes from whole-genome shotgun short reads. Genome Research, 24 (8), 1384–1395.

Kayvanfar, N., Aliabadian, M., Niu, X., Zhang, Z., & Liu, Y. (2017). Phylogeography of the Common Pheasant Phasianus colchicus. Ibis, 159 (2), 430–442.

Liu, N., & Sun, H. (1992). The geographic variance and classification of Phasianus colchicus strauchi. J. Lanzhou Univ, 28 (2), 135–139.

Liu, Y., Zhan, X., Wang, N., Chang, J., & Zhang, Z. (2010). Effect of geological vicariance on mitochondrial DNA differentiation in Common Pheasant populations of the Loess Plateau and eastern China. Molecular Phylogenetics and Evolution, 55 (2), 409–417.

Lomsadze, A., Ter-Hovhannisyan, V., Chernoff, Y. O., & Borodovsky, M. (2005). Gene identification in novel eukaryotic genomes by self-training algorithm. Nucleic acids research, 33 (20), 6494–6506.

Long, S., Zhou, C.-q., Wang, W.-k., Wei, W., & Hu, J.-c. (2007). The Habitat and Nestsite Selection of Common Pheasants in Spring and Summer in Nanchong, China. Zoological Research, 28 (3), 249.

Madden, T. (2013). The BLAST sequence analysis tool.

Madge, S., McGowan, P. J., & Kirwan, G. M. (2002). Pheasants, partridges and grouse: a guide to the pheasants, partridges, quails, grouse, guineafowl, buttonquails and sandgrouse of the world: A&C Black.

Marçais, G., & Kingsford, C. (2011). A fast, lock-free approach for efficient parallel counting of occurrences of k-mers. Bioinformatics, 27 (6), 764–770.

Miller, M. R., Dunham, J. P., Amores, A., Cresko, W. A., & Johnson, E. A. (2007). Rapid and cost-effective polymorphism identification and genotyping using restriction site associated DNA (RAD) markers. Genome Research, 17 (2), 240–248.

Nadachowska-Brzyska, K., Burri, R., Olason, P. I., Kawakami, T., Smeds, L., & Ellegren, H. (2013). Demographic divergence history of pied flycatcher and collared flycatcher inferred from whole-genome re-sequencing data. PLoS Genetics, 9 (11), e1003942.

Nosil, P., Funk, D. J., & ORTIZ-BARRIENTOS, D. (2009). Divergent selection and heterogeneous genomic divergence. Molecular Ecology, 18 (3), 375–402.

Pfarr, J. (2012). True Pheasants: a noble quarry: Hancock House Publishers.

Poelstra, J. W., Vijay, N., Bossu, C. M., Lantz, H., Ryll, B., Müller, I., … Grabherr, M. G. (2014). The genomic landscape underlying phenotypic integrity in the face of gene flow in crows. Science, 344 (6190), 1410–1414.

Price, A. L., Jones, N. C., & Pevzner, P. A. (2005). De novo identification of repeat families in large genomes. Bioinformatics, 21 (suppl_1), i351–i358.

Qu, J., Liu, N., Bao, X., & Wang, X. (2009). Phylogeography of the ring-necked pheasant (Phasianus colchicus) in China. Molecular Phylogenetics and Evolution, 52 (1), 125–132.

Robertson, P. A. (1996). Does nesting cover limit abundance of ring-necked pheasants in North America? Wildlife Society Bulletin, 98–106.

Salamov, A. A., & Solovyev, V. V. (2000). Ab initio gene finding in Drosophila genomic DNA. Genome Research, 10 (4), 516–522.

Simão, F. A., Waterhouse, R. M., Ioannidis, P., Kriventseva, E. V., & Zdobnov, E. M. (2015). BUSCO: assessing genome assembly and annotation completeness with single-copy orthologs. Bioinformatics, 31 (19), 3210–3212.

Slater, G. S. C., & Birney, E. (2005). Automated generation of heuristics for biological sequence comparison. BMC Bioinformatics, 6 (1), 31.

Smit, A., & Hubley, R. (2008). RepeatModeler Open-1.0. http://www.repeatmasker.org.

Smit, A., Hubley, R., & Green, P. (2016). RepeatMasker Open-4.0. http://www.repeatmasker.org.

Stanke, M., & Waack, S. (2003). Gene prediction with a hidden Markov model and a new intron submodel. Bioinformatics, 19 (suppl_2), ii215–ii225.

Toews, D. P., Taylor, S. A., Vallender, R., Brelsford, A., Butcher, B. G., Messer, P. W., & Lovette, I. J. (2016). Plumage genes and little else distinguish the genomes of hybridizing warblers. Current Biology, 26 (17), 2313–2318.

Turner, T. L., Hahn, M. W., & Nuzhdin, S. V. (2005). Genomic islands of speciation in Anopheles gambiae. PLoS Biology, 3 (9), e285.

Wang, B., Xie, X., Liu, S., Wang, X., Pang, H., & Liu, Y. (2017). Development and characterization of novel microsatellite markers for the Common Pheasant (Phasianus colchicus) using RAD-seq. Avian Research, 8 (1), 4.

Wu, C. I. (2001). The genic view of the process of speciation. Journal of Evolutionary Biology, 14 (6), 851–865.

Zhang, L., An, B., Backström, N., & Liu, N. (2014). Phylogeography-based delimitation of subspecies boundaries in the Common Pheasant (Phasianus colchicus). Biochemical Genetics, 52 (1–2), 38–51.

