## Supplementary information for "Genome assembly of the common pheasant *Phasianus colchicus*, a model for speciation and ecological genomics"

**Appendix Population genomic analysis using RAD-sequencing**

**To facilitate future population genomics studies of the common pheasant, we performed restriction site-associated DNA sequencing (RAD sequencing) (Miller et al. 2007) for an additional 45 male individuals from seven subspecies across China (Figure 3a, Table S4). Genomic DNAs were digested with a restriction enzyme, ApeKI. Then the sequencing libraries were prepared following the manufacturer’s protocol and sequenced on Illumina HiSeq 2000 platform. About 127.98G bases were generated (raw data) for all the samples. The raw data were processed in two steps: deleting adapter sequences and removing problematic reads (when the rate of low quality (quality value <=5 (E) was more than or equal to 50%). About 123.27 G bases were generated (clean data) in this step. Then all reads were assigned to the individuals by the unique barcodes and the specific recognition site using Sabre (github.com/najoshi/sabre). The reads without the unique barcodes and the specific sequences were discarded. Final ‘reads 1’ length was trimmed to 82 nucleotides (minimal length). After this step, about 114.91 Gb were generated (clean data) for all 45 samples (Table S4).**

**SNP calling**

**The well-assembled common pheasant genome was used as reference genome that allows for mapping of the RAD sequencing reads using BWA v0.7.15 (Li & Durbin 2009). SNPs were extracted from mapped reads using ‘pileup’ in SAMtools v1.4 with default options (Li et al. 2009, Li 2011), and also with Genome Analysis Toolkit (GATK 3.7) with parameters "QD < 2.0 || FS > 60.0 || MQ < 40.0 || MQRankSum < -12.5 || ReadPosRankSum < -8.0" (McKenna et al. 2010).**

**Nucleotide differences called by SAMtools were classified as SNPs if (i) the alternate allele was supported by a minimum coverage of 3, and (ii) the alternate allele had a minimum Phred quality score of 20. Nucleotide differences called by GATK were classified as SNPs if they were flagged as “PASS”. We used SnpEff (version 4.3) (Cingolani et al.2012) to identify the fine-scale distribution of SNPs.**

**To infer population structuring of the common pheasant in northern China, we carried out a genetic admixture analysis based on the obtained SNP dataset using ADMIXTURE 1.3 (Alexander et al. 2009). This approach applies model-based estimation of ancestry in unrelated individuals from their SNP genotype datasets. We defined K, the potential number of ancestral genetic cluster and ranged K from 1 to 8 with 120 iterations at each K. In addition, we conducted a model-free multivariate method, i.e. principal component analysis (PCA)** using the smartpca program of the EIGENSOFT package (Price et al. 2006)**. This method analyzes a genetic covariance matrix calculated from genotypes, and we visualized the PCA results based on the variance explained by the first two axes.**

**Literature Cited**

**-Alexander DH, Novembre J and Lange K. Fast model-based estimation of ancestry in unrelated individuals. Genome Research. 2009;19 9:1655-64.**

**-Cingolani P, Platts A, Wang LL, Coon M, Nguyen T, Wang L, et al. A program for annotating and predicting the effects of single nucleotide polymorphisms, SnpEff: SNPs in the genome of Drosophila melanogaster strain w1118; iso-2; iso-3. Fly. 2012;6 2:80-92.**

**-Li H and Durbin R. Fast and accurate short read alignment with Burrows–Wheeler transform. Bioinformatics. 2009;25 14:1754-60.**

**-Li H, Handsaker B, Wysoker A, Fennell T, Ruan J, Homer N, et al. The sequence alignment/map format and SAMtools. Bioinformatics. 2009;25 16:2078-9.**

**-Li H. A statistical framework for SNP calling, mutation discovery, association mapping and population genetical parameter estimation from sequencing data. Bioinformatics. 2011;27 21:2987-93.**

**-McKenna A, Hanna M, Banks E, Sivachenko A, Cibulskis K, Kernytsky A, et al. The Genome Analysis Toolkit: a MapReduce framework for analyzing next-generation DNA sequencing data. Genome Research. 2010;20 9:1297-303.**

**-Miller MR, Dunham JP, Amores A, Cresko WA and Johnson EA. Rapid and cost-effective polymorphism identification and genotyping using restriction site associated DNA (RAD) markers. Genome Research. 2007;17 2:240-8.**

**-Price A L, Patterson N J, Plenge R M, et al. Principal components analysis corrects for stratification in genome-wide association studies. Nature Genetics, 2006; 38(8): 904-9.**

**Table S1.** Summary statistics of the generated whole genome shotgun sequencing data

| Library name | Insert size | Raw paired reads | Raw base (bp) | Filtered paired reads | Filtered base (bp) | Sequence depth(x) |
| --- | --- | --- | --- | --- | --- | --- |
| DES00955_L1 | 250bp | 63000115 | 18900034500 | 56271403 | 16881420900 | 16.38 |
| DES00954_L1 | 250bp | 60206332 | 18061899600 | 53867022 | 16160106600 | 15.68 |
| DES00953_L1 | 450bp | 55935789 | 16780736700 | 51867389 | 15560216700 | 15.10 |
| DES00952_L1 | 450bp | 59938896 | 17981668800 | 55938980 | 16781694000 | 16.28 |
| DEL00730_L4 | 2kbp | 51643562 | 15493068600 | 44719493 | 13415847900 | 13.02 |
| DEL00731_L4 | 5kbp | 61115241 | 18334572300 | 32411208 | 9723362400 | 9.44 |
| DEL00732_L4 | 10kbp | 38687097 | 11606129100 | 21205997 | 6361799100 | 6.17 |

**Table S2. Summary** statistics of the genome assembly for the common pheasant.

| Assembled Common Pheasant Genome | |
| --- | --- |
| Total length | 1,023,273,632 bp |
| Number of scaffolds | 82,771 |
| N50 of contigs | 178,013 bp |
| N50 of scaffolds | 10,186,719 bp |
| Longest scaffolds | 42,030,034 bp |
| Average scaffold length | 12,362.71 bp |
| GC level | 41.24% |

**Table S3. Summary of BUSCO analysis for the common pheasant.**

| Types of BUSCOs | Count | Ratio |
| --- | --- | --- |
| Complete BUSCOs | 4585 | 93.3% |
| Complete and single-copy BUSCOs | 4532 | 92.2% |
| Complete and duplicated BUSCOs | 53 | 1.1% |
| Fragmented BUSCOs | 205 | 4.2% |
| Missing BUSCOs | 125 | 2.5% |

Table S4. Summary of the RAD sequencing data and SNP detection in populations of seven subspecies of the common pheasant.

| **Population** | **ID** | **Location** | **Reads (M)** | | **Bases (Mb)** | **GC (%)** | **SNPs** |
| --- | --- | --- | --- | --- | --- | --- | --- |
| *shawi* | 1 | Shache, Xinjiang | | 21.01 | 1817.7 | 39.14 | 741515 |
|  | 2 | Shache, Xinjiang | | 23.01 | 1979 | 39.2 | 728400 |
|  | 3 | Wensu, Xinjiang | | 12.69 | 1104.1 | 39.16 | 624846 |
|  | 4 | Wensu, Xinjiang | | 23.01 | 2002.1 | 39.14 | 749513 |
|  | 5 | Wensu, Xinjiang | | 21.85 | 1922.5 | 38.98 | 774408 |
|  | 6 | Wensu, Xinjiang | | 17.36 | 1518.6 | 39.2 | 683611 |
|  | 7 | Wensu, Xinjiang | | 23.43 | 2050.4 | 39.12 | 782767 |
| *vlangalii* | 8 | Golmud, Qinghai | | 34.25 | 2980 | 39.01 | 792904 |
|  | 9 | Golmud, Qinghai | | 19.71 | 1704.5 | 38.89 | 689226 |
|  | 10 | Golmud, Qinghai | | 24.47 | 2116.5 | 39.03 | 613075 |
|  | 11 | Golmud, Qinghai | | 26.3 | 2261.5 | 39.24 | 711822 |
|  | 12 | Golmud, Qinghai | | 32.4 | 2786.5 | 38.81 | 733846 |
|  | 13 | Qaidam, Qinghai | | 21.65 | 1861.9 | 38.97 | 616743 |
|  | 14 | Qaidam, Qinghai | | 21 | 1826.6 | 39.08 | 708986 |
| *strauchi* | 15 | Xining, Qinghai | | 30.57 | 2659.4 | 38.93 | 869430 |
|  | 16 | Xining, Qinghai | | 25.59 | 2252.1 | 38.71 | 836985 |
|  | 17 | Xining, Qinghai | | 29.16 | 2552 | 39.3 | 916936 |
|  | 18 | Xining, Qinghai | | 18.06 | 1589.7 | 39.15 | 734053 |
|  | 19 | Datong, Qinghai | | 20.74 | 1793.8 | 39.15 | 797247 |
|  | 20 | Datong, Qinghai | | 14.91 | 1289.5 | 39.02 | 707193 |
| *kiansuensis* | 21 | Ningwu, Shanxi | | 21.83 | 1877.7 | 38.91 | 717772 |
|  | 22 | Ningwu, Shanxi | | 27.2 | 2339.1 | 39.24 | 828957 |
|  | 23 | Hohhot, Inner Mongolia | | 12.27 | 1067.7 | 39.14 | 615167 |
|  | 24 | Hohhot, Inner Mongolia | | 25.34 | 2223 | 39.37 | 926647 |
| *karpowi* | 25 | Chengde, Hebei | | 15.76 | 1371.1 | 38.97 | 762361 |
|  | 26 | Chengde, Hebei | | 26.55 | 2296.9 | 38.7 | 936224 |
|  | 27 | Chengde, Hebei | | 19.75 | 1707.9 | 38.99 | 823696 |
|  | 28 | Chifeng, Inner Mongolia | | 29.7 | 2554.1 | 39.04 | 899788 |
|  | 29 | Chifeng, Inner Mongolia | | 24.12 | 2074.5 | 39.06 | 841414 |
|  | 30 | Chifeng, Inner Mongolia | | 20.57 | 1768.8 | 39.02 | 821887 |
|  | 31 | Chifeng, Inner Mongolia | | 19.24 | 1654.5 | 39.33 | 740318 |
|  | 32 | Chifeng, Inner Mongolia | | 16.47 | 1416.5 | 38.88 | 763730 |
| *torquatus* | 33 | Zhenjiang, Jiangsu | | 15.13 | 1316.7 | 38.82 | 734093 |
|  | 34 | Zhenjiang, Jiangsu | | 23.29 | 2026.6 | 38.95 | 864847 |
|  | 35 | Zhenjiang, Jiangsu | | 27.96 | 2460.5 | 39 | 999733 |
|  | 36 | Jiangshan, Zhejiang | | 21.26 | 1839.1 | 39.12 | 849350 |
|  | 37 | Xinyang, Henan | | 19.71 | 1694.7 | 39.34 | 818415 |
|  | 38 | Xinyang, Henan | | 16.22 | 1394.8 | 39.05 | 762824 |
|  | 39 | Xinyang, Henan | | 24.41 | 2099.3 | 39.24 | 879765 |
|  | 40 | Pengze, Jiangxi | | 16.63 | 1429.9 | 38.93 | 784304 |
| *elegans* | 41 | Shangri-La, Yunnan | | 14.29 | 1243.6 | 38.84 | 734041 |
|  | 42 | Yuxi, Yunnan | | 36.57 | 3209 | 39.2 | 924150 |
|  | 43 | Yuxi, Yunnan | | 23.02 | 1991.3 | 39.14 | 820069 |
|  | 44 | Dali, Yunna | | 14.24 | 1239.2 | 39.24 | 710531 |
|  | 45 | Dali, Yunna | | 23.73 | 2078.1 | 38.87 | 328473 |

Table S5. The distribution of the SNPs in the common pheasant genome.

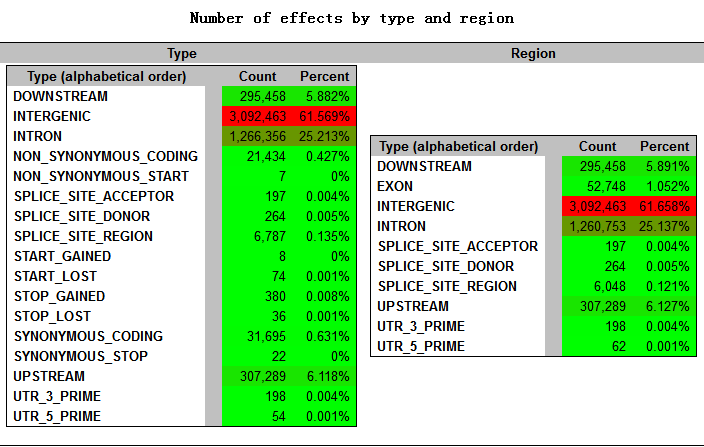

Figure S1. 19-mer distribution of the common pheasant genome using Jellyfish with all paired-end sequencing data.

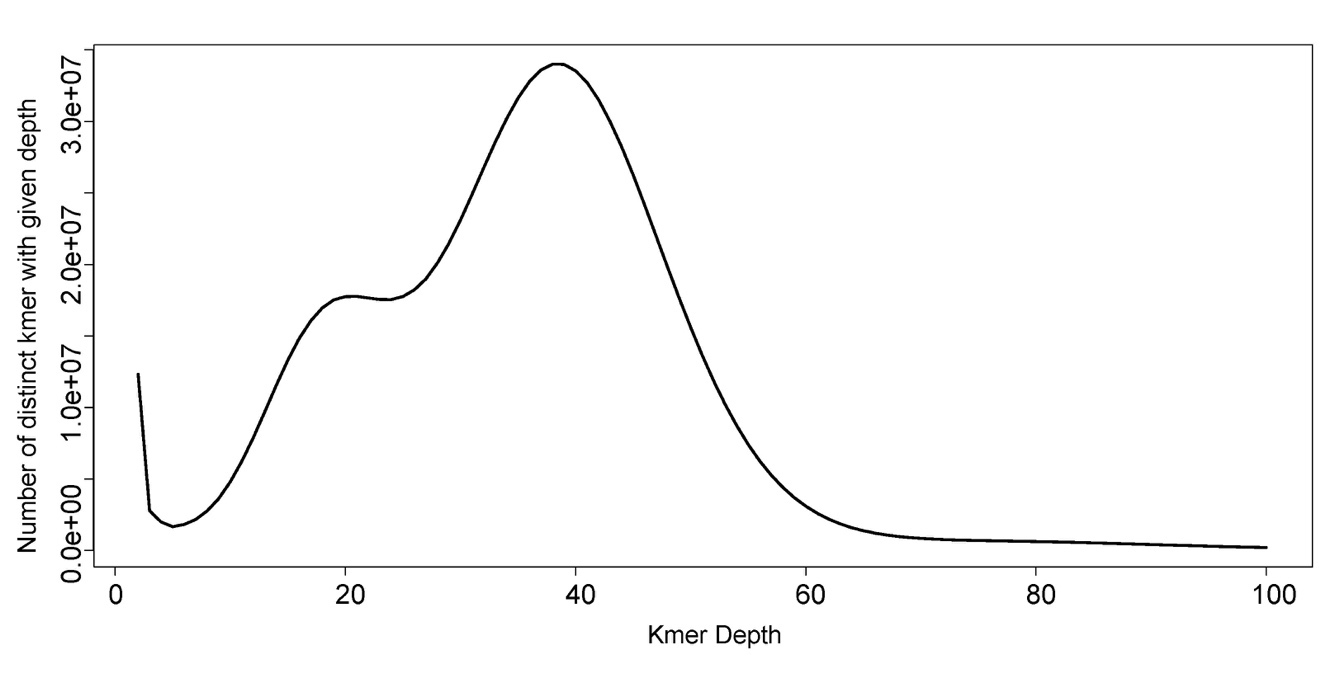

Figure S2. Shown is the cross-validation plot for different numbers of genetic groups (K), in which four genetic cluster is inferred since the cross-validation error has the lowest value among the alternatives.

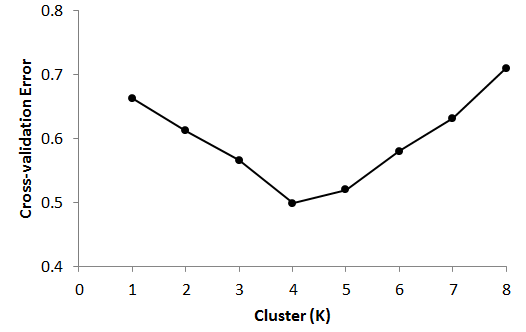

Figure S3. Principal component analysis (PCA) of 45 individuals from seven subspecies of common pheasant using 4,376,351 SNPs. Colour dots represent different subspecies.

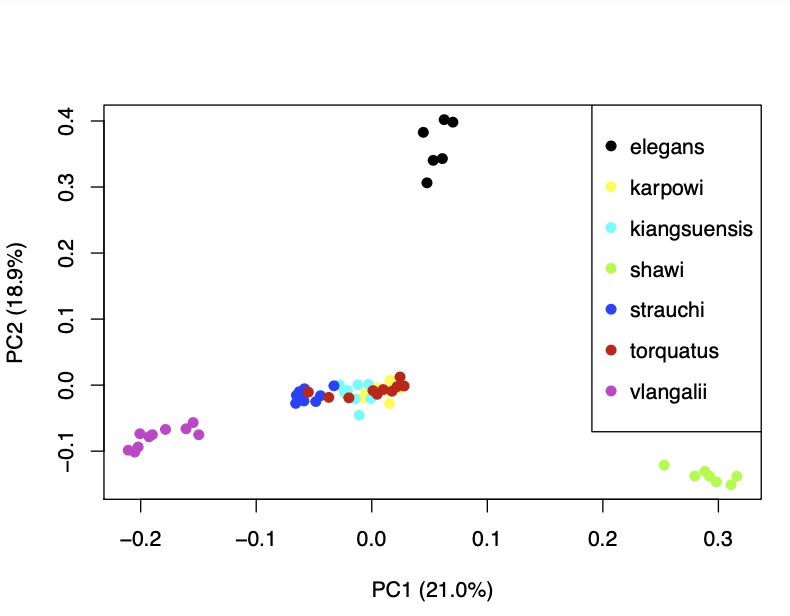
